# c-Myc Represses Transcription of the Epstein-Barr Virus Latent Membrane Protein 1 Early After Primary B Cell Infection

**DOI:** 10.1101/168484

**Authors:** Alexander M. Price, Joshua E. Messinger, Micah A. Luftig

**Author notes:** Present address: Pathology and Laboratory Medicine, The Children’s Hospital of Philadelphia (Alexander M. Price). These authors contributed equally to this work (Alexander M. Price, Joshua E. Messinger). Corresponding author: Micah Luftig.

## Abstract

Recent evidence has shown that the EBV oncogene LMP1 is not expressed at high levels early after EBV-infection of primary B cells, despite its being essential for the long-term outgrowth of immortalized lymphoblastoid cell lines (LCLs). In this study, we found that expression of LMP1 increased fifty-fold between seven days post infection and the LCL state. Metabolic labeling of nascently transcribed mRNA indicated this was primarily a transcription-mediated event. EBNA2, the key viral transcription factor regulating LMP1, and CTCF, an important chromatin insulator, were recruited to the LMP1 locus similarly early and late after infection. However, the activating histone H3K9Ac mark was enriched at the LMP1 promoter in LCLs relative to early-infected B cells. We found that high c-Myc activity in EBV-infected lymphoma cells as well as overexpression of c-Myc in an LCL model system repressed LMP1 transcription. Finally, we found that chemical inhibition of c-Myc expression both in LCLs and early after primary B-cell infection increased LMP1 expression. These data support a model in which high levels of endogenous c-Myc activity induced early after primary B-cell infection directly represses LMP1 transcription.

**IMPORTANCE:** EBV is a highly successful pathogen that latently infects greater than 90% of adults worldwide and is also causally associated with a number of B-cell malignancies. EBV expresses a set of viral oncoproteins and non-coding RNAs during the latent life cycle with the potential to promote cancer. Critical among these is the viral latent membrane protein, LMP1. Prior work suggests that LMP1 is essential for EBV to immortalize B cells, but our recent work indicates that LMP1 is not produced at high levels during the first few weeks after infection. Here, we show that the transcription of LMP1 can be negatively regulated by a host transcription factor, c-Myc. Ultimately, understanding the regulation of EBV-encoded oncogenes will allow us to better treat cancers that rely on these viral products for survival.

## INTRODUCTION

Epstein-Barr virus (EBV) infection of primary human B cells leads to their immortalization, or growth transformation (1). These immortalized cells, called lymphoblastoid cell lines (LCLs), are latently infected by EBV and express a program, called latency III, consisting of nine viral proteins and many non-coding RNAs. LCLs expressing Latency III genes are a model system for EBV-associated malignancies such as Post-Transplant Lymphoproliferative Disease (PTLD) and Diffuse Large B Cell Lymphoma (DLBCL) (1). Characteristic of Latency III is expression of the essential viral oncogene latent membrane protein 1 (LMP1), which is a constitutively active TNF receptor homologue that induces the host NFκB and AP-1 signaling pathways to promote survival (2-5). LMP1 is both necessary for EBV immortalization of primary human B cells and sufficient to induce lymphomas when expressed in the murine B cell compartment (6, 7). The regions of the LMP1 cytoplasmic tail that engage TNFR associated factors (TRAFs) and TNFR associated death domain (TRADD) responsible for NFκB activation are coincident with the regions required for EBV-mediated immortalization (8, 9). LCLs are functionally addicted to the NFκB signaling induced by LMP1 as inhibition of this pathway results in apoptosis (3).

Studies of LMP1 transcriptional regulation have defined cis-acting elements and transacting factors important in controlling its expression. In epithelial cells, LMP1 expression is driven by a unique promoter found within the viral terminal repeats (10, 11). However, transcription of LMP1 in Latency III expressing B cells is initiated at a bi-directional LMP1/LMP2B promoter (12). The critical activator of this Latency III promoter is Epstein-Barr Nuclear Antigen 2 (EBNA2), which binds to a so-called EBNA2-response element (E2RE) through the host factors RBP-Jκand PU.1 (13, 14). Other viral factors, such as EBNA3C, have also been shown to aid in the transcription of LMP1 (15, 16). Additionally, a number of host factors have been shown to fine-tune LMP1 transcription, including ATF4 (17, 18), IRF7 (19), and NFκB subunits themselves (20, 21). In fact, host factors often lead to auto-regulatory feedback loops that maintain the levels of LMP1 expression, whether it is NFκB in B cells (20) or STAT signaling on the terminal repeat promoter in epithelial cells (22). Furthermore, the epigenetic state of the LMP1 promoter plays a key role in transcription. Changes in cell type and latency state alter the prevalence of active and repressive histone marks on the LMP1 promoter as well as CpG DNA methylation (23). In addition, viral chromatin architecture and enhancer looping mediated by host factors such as CTCF and Rad21 play a direct role in the full activation of LMP1 (24).

LMP1 mRNA is also post-transcriptionally regulated to control protein output. While LMP1 is one of the most abundant viral transcripts detected in latently infected cells, it is transcribed at a much lower rate implying a certain level of mRNA stability (25). This stability can be counteracted by micro-RNA (miRNA) targeting of the long LMP1 3’ untranslated region. LMP1 can be targeted by both EBV-encoded viral miRNAs such as miR-BART1-5p, miR-BART16-3p, and miR-BART17-5p (26) as well as a number of host miRNAs (27). Of these host miRNAs, inhibiting the c-Myc regulated miR17~92 miRNA cluster led to an upregulation of LMP1 protein levels and slowed cell growth (27). This slowed growth could be explained in part by the known ability of overexpressed LMP1 to be cytostatic (28-30).

The cellular oncogene c-Myc is overexpressed in a wide variety of cancers, including EBV-associated Burkitt lymphoma where c-Myc is expressed at high levels due to a characteristic chromosomal 8:14 translocation. Additionally, EBV induces c-Myc expression upon infection of B cells by utilizing EBNA2 to co-opt native super-enhancer architecture upstream of c-Myc (31). Upon *de novo* infection, c-Myc levels peak within the first week after infection, then wane, but remain elevated throughout LCL outgrowth where c-Myc is crucial for the maintenance of the LCL phenotype (32, 33). This conflicts with the NFκB addiction observed in LCLs, as the c-Myc and NFκB signaling pathways have been shown to be directly incompatible (34).

We have previously shown that, despite being detectable early after infection of primary B cells, LMP1 does not reach LCL levels of expression until greater than two weeks post infection (35). These EBV-infected cells express EBNA2, proliferate, and show no signs of apoptosis even when NFκB is inhibited during the first weeks of infection (35, 36). Furthermore, the exact role of LMP1 in the EBV life cycle has been called into question by recent work showing that LMP1 is dispensable for tumor formation in a humanized mouse model (37, 38). In this work, we address the question of how LMP1 is delayed in expression from early to late times after primary B-cell infection. We investigate the nature of the mRNA change and the role of cellular factors in temporal regulation of LMP1 expression.

## RESULTS

### LMP1 transcription is robustly increased from early to late times after primary B cell infection

LMP1 mRNA and protein levels and, consequently, NFκB targets are significantly lower during the first two weeks following EBV infection of primary B cells relative to that found in immortalized lymphoblastoid cell lines (LCLs) (35). Prior studies indicate that the LMP1 mRNA is the most abundant latency transcript in LCLs despite being poorly transcribed (25). Therefore, to determine the mechanism for low LMP1 mRNA levels at early times after infection, we assayed the relative transcription rate and half-life using a 4-thiouridine (4sU) metabolic labeling approach (**Fig. 1A** and (39)). This was performed by using Fluorescence Activated Cell Sorting (FACS) to generate a pure population of proliferating (Celltrace Violet^lo^) EBV-infected B cells six days after primary human Peripheral Blood Mononuclear Cell (PBMC) infection and allowing the cells to rest overnight. Twenty-four hours later (Day 7 post infection), cells were pulsed with 4sU for exactly one hour before total RNA was harvested. The process was repeated for LCLs that grew out from matched PBMC donors five weeks post infection. As previously observed, the total LMP1 mRNA level increased ~50-fold from day 7 post infection through LCL outgrowth (**Fig. 1B**). Comparing the ratio of 4sU labeled nascent RNA and unlabeled decaying RNA (39), we found that the halflife of the LMP1 mRNA increased two-fold between early and late times after infection from ~2 hours at 7 days post infection to ~4 hours in LCLs (**Fig. 1C**). Over the same time frame the relative transcription rate of LMP1 mRNA increased nearly 25-fold (**Fig. 1D**). We also queried the EBNA2-specific mRNA as well as Cp-derived EBNA transcripts and found only modest differences in their overall expression, transcription rate, or stability through B cell outgrowth (**Fig. 1B-D**).

**Figure 1.**
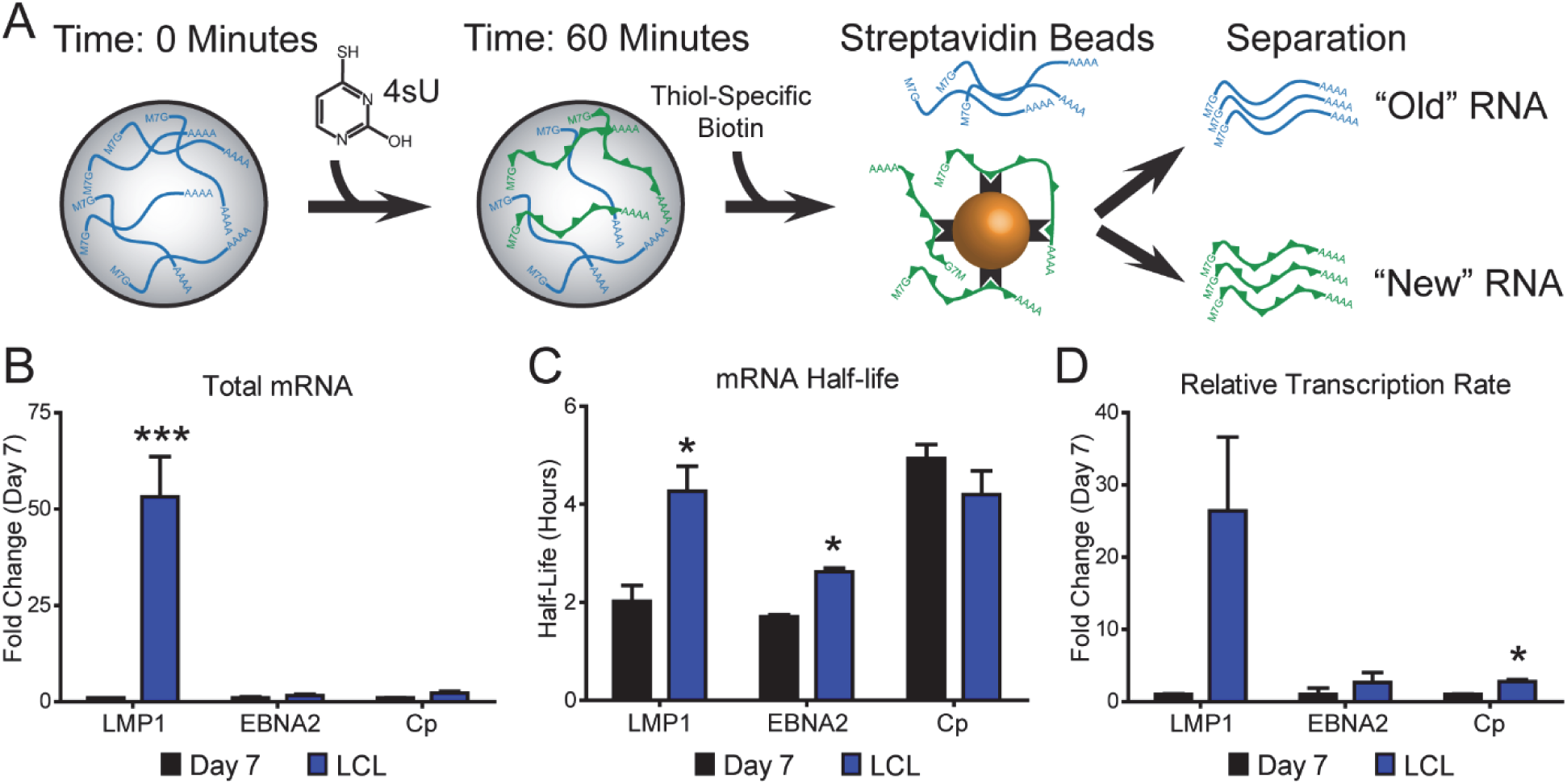
4sU metabolic pulsing reveals differences in LMP1 mRNA abundance, half-life and transcription rate at Day 7 compared to LCL. (A) Schematic describing 4sU experiments. Cells are pulsed for 1 hour with 4sU before RNA isolation by Trizol. 4sU labeled RNA is then conjugated to biotin and isolated via magnetic streptavidin. Unbound RNA is saved as the “old” fraction while bound RNA eluted from the beads becomes the “new” fraction. Isolated RNA is then reverse transcribed for RT-qPCR analysis. (B) RT-qPCR analysis of total mRNA from 4sU experiments for indicated genes at indicated times post infection. (C) RT-qPCR analysis of mRNA half-life from 4sU experiments. Half-life is normalized to GAPDH. (D) Transcription rate analysis of mRNA from 4sU experiments for indicated genes at indicated times post infection. All transcription rates are relative to GAPDH. For all panels, Cp denotes EBV C promoter transcripts. Each bar is representative of at least 3 independent donors. Error bars represent SEM. ^∗^ denotes p<0.05, ^∗∗^ denotes p<0.01 and ^∗∗∗^ denotes p<0.001 by one-tailed pairwise student’s t-test.

### The LMP1 locus is occupied by EBNA2 and CTCF similarly early and late after infection; however lower levels of activated histones are detected early

In B cells, the transcription of LMP1 is regulated by EBNA2 (40). Prior work indicates that EBNA2-regulated genes, including c-Myc and CD23, are induced to higher levels at day 7 post infection than in LCLs suggesting that EBNA2 activity is not broadly suppressed during early infection (32). However, it remained a possibility that LMP1 expression was low during early infection because of poor EBNA2 recruitment to the bi-directional LMP1/2B promoter. To address this, we performed Chromatin Immuno-precipitation (ChIP) for EBNA2 at day 7 post infection and in LCLs. LMP1 and LMP2 are transcribed from a complex and overlapping region of the EBV episome near the terminal repeats (TR), diagrammed in **Fig. 2A**. We found that EBNA2 was recruited comparably to the LMP1 promoter and the viral C promoter both early after infection and in LCLs (**Fig. 2B**). A region in the gene body of EBNA3C distal to both the LMP1p and the Cp was used as a negative control (**Fig. 2B**). Similarly, EBNA2 was recruited to the two major c-Myc upstream enhancer loci (31) as well as the CD23 promoter comparably at 7 days post infection as in LCLs (**Fig. 2C**). Despite the lack of change in EBNA2 occupancy at early and late times post infection, the activating histone mark H3K9 Acetyl was enriched at the LMP1p in LCLs, correlating with the observed increased level of transcription (**Fig. 2D**). A recent publication has implicated the chromatin architecture mediated by CTCF at a single CTCF site downstream of LMP1 (here called the CTCF-Response Element) to be important for H3K9 Acetylation and transcription at the LMP1 promoter (24). We assayed for CTCF-occupancy at both this CTCF-RE as well as the EBV enhancer found in the EBER/OriP region of the genome and found that CTCF occupancy did not change between day 7 post infection cells and LCLs (**Fig. 2E**).

**Figure 2.**
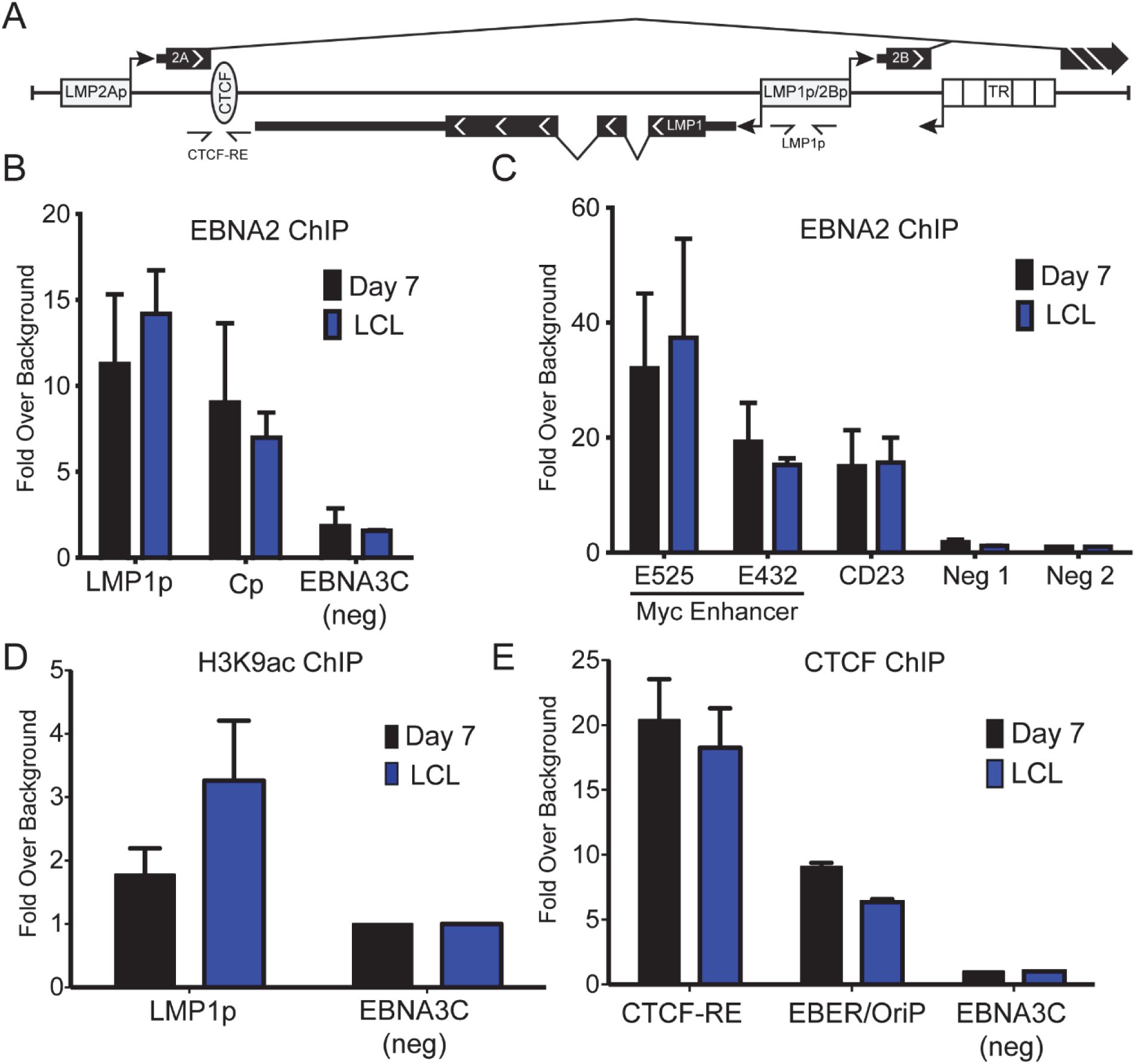
Interrogation of the LMP1 locus by ChIP-PCR. (A) Schematic of the LMP1 genomic locus. Arrows with half arrow heads denote the location of primers used for ChIP-PCR in (B), (D) and (E). (B) Sorted Day 7 proliferating primary EBV-infected B cells (black) or fully outgrown (>35 days) LCLs (blue) were harvested for chromatin and ChIP for EBNA2 was performed. LMP1p denotes the LMP1 promoter, Cp denotes the C promoter and EBNA3C is an EBV genomic negative control. All samples are normalized to the average of the two negative controls from the human genome in (C). (C) Same experiment as in (B) but RT-qPCR was performed for regions of the human genome. Neg1 and Neg2 denote transcription factor deserts within the human genome serving as a negative control. All samples are normalized to the average of the two negative controls. (D) Similar experiment as in (B) but ChIP was performed for H3K9ac. All samples are normalized to EBNA3C. (E) Similar experiments as in (B) but ChIP was performed for CTCF. CTCF-RE denotes the CTCF binding site in the LMP1 locus (panel A), EBER/OriP denotes a known CTCF binding site by EBER/OriP. For all panels, all bars represent the average of at least 3 independent donors. All error bars represent SEM.

### c-Myc suppresses LMP1 transcription

Despite EBNA2 recruitment to the LMP1 promoter, reduced H3K9 acetylation suggested poor LMP1 transcription. Early after infection when LMP1 expression is lowest, c-Myc is strongly activated by EBNA2, and during the time frame that LMP1 expression increases the expression of c-Myc and its targets are subsequently attenuated (32). Given this correlation, we sought to test the role of c-Myc in the regulation of LMP1 expression. We first assayed LMP1 transcription using 4sU labeling in EBV-infected Burkitt lymphoma (BL) cell lines that express very high levels of c-Myc relative to LCLs. We found that LMP1 mRNA and protein levels were higher in LCLs than in EBV-infected BL41 cells (**Fig. 3A-B**). Consistent with our hypothesis, the transcription of LMP1 was significantly lower in EBV-infected BL41 cells relative to LCLs (**Fig. 3C**).

**Figure 3.**
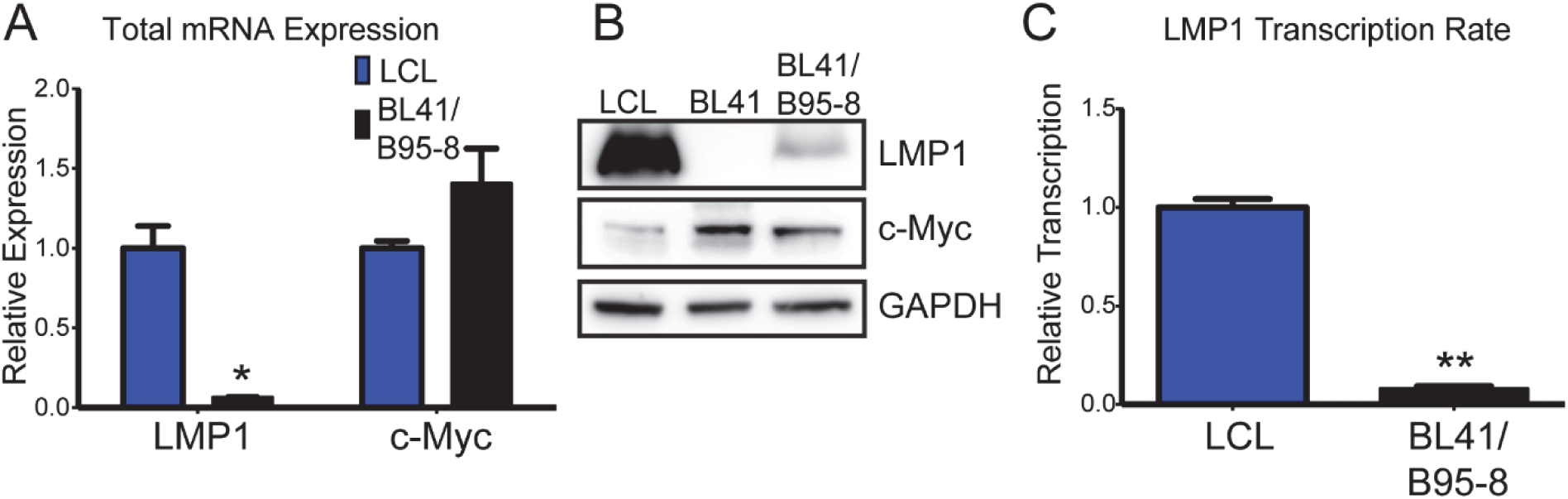
Increased c-Myc in BL41/B95-8 correlates with decreased LMP1 mRNA abundance, transcription rate and protein. (A) RT-qPCR analysis for total mRNA from fully immortalized (>35 days) LCLs or BL41/B95-8 for indicated genes. All values are normalized to SETDB1. (B) Western blot analysis for indicated proteins in LCL, EBV negative BL41 and BL41 re-infected with EBV, BL41/B95-8. (C) Relative transcription rate from 4sU metabolic labeling of LMP1 mRNA transcription rate in LCL and BL41/B95-8. All values are relative to GAPDH. For each panel, each bar is representative of 3 independent donors/experiments. All error bars represent SEM. ^∗^ denotes p<0.05 and ^∗∗^ denotes p<0.01 by one tailed student’s pairwise t-test.

To directly assess the effect of c-Myc on LMP1 transcription in an LCL, we used the P493-6 model of c-Myc and EBNA2 regulation (41, 42). In this system, EBNA2 (encoded endogenously from the viral genome) is controlled post-translationally by Estrogen (β-Estradiol) due to its fusion to a modified estrogen receptor, and heterologous c-Myc expression is provided *in trans* and controlled transcriptionally through a tetracycline (tet-off) system. We verified that induction of EBNA2 and inactivation of c-Myc (Estrogen+, Tetracycline+) led to an LCL-like phenotype with high levels of LMP1 transcription (**Fig. 4A**). However, when removing EBNA2 activity (Estrogen-, Tetracycline-) or inducing EBNA2 in the presence of high levels of c-Myc (Estrogen+, Tetracycline-), we observed blunted endogenous LMP1 transcription relative to that induced in the presence of low levels of c-Myc (**Fig. 4A**). These changes in transcription correlated with the levels of total LMP1 mRNA and protein (**Fig. 4B-C**), while EBNA2-ER transcription from the endogenous Cp was not affected by c-Myc overexpression (**Fig. 4B**). This controlled experiment indicated that c-Myc overexpression is sufficient to suppress LMP1 transcription in an LCL and suggested that c-Myc may be critical for suppression of LMP1 transcription early after primary B-cell infection as well.

**Figure 4.**
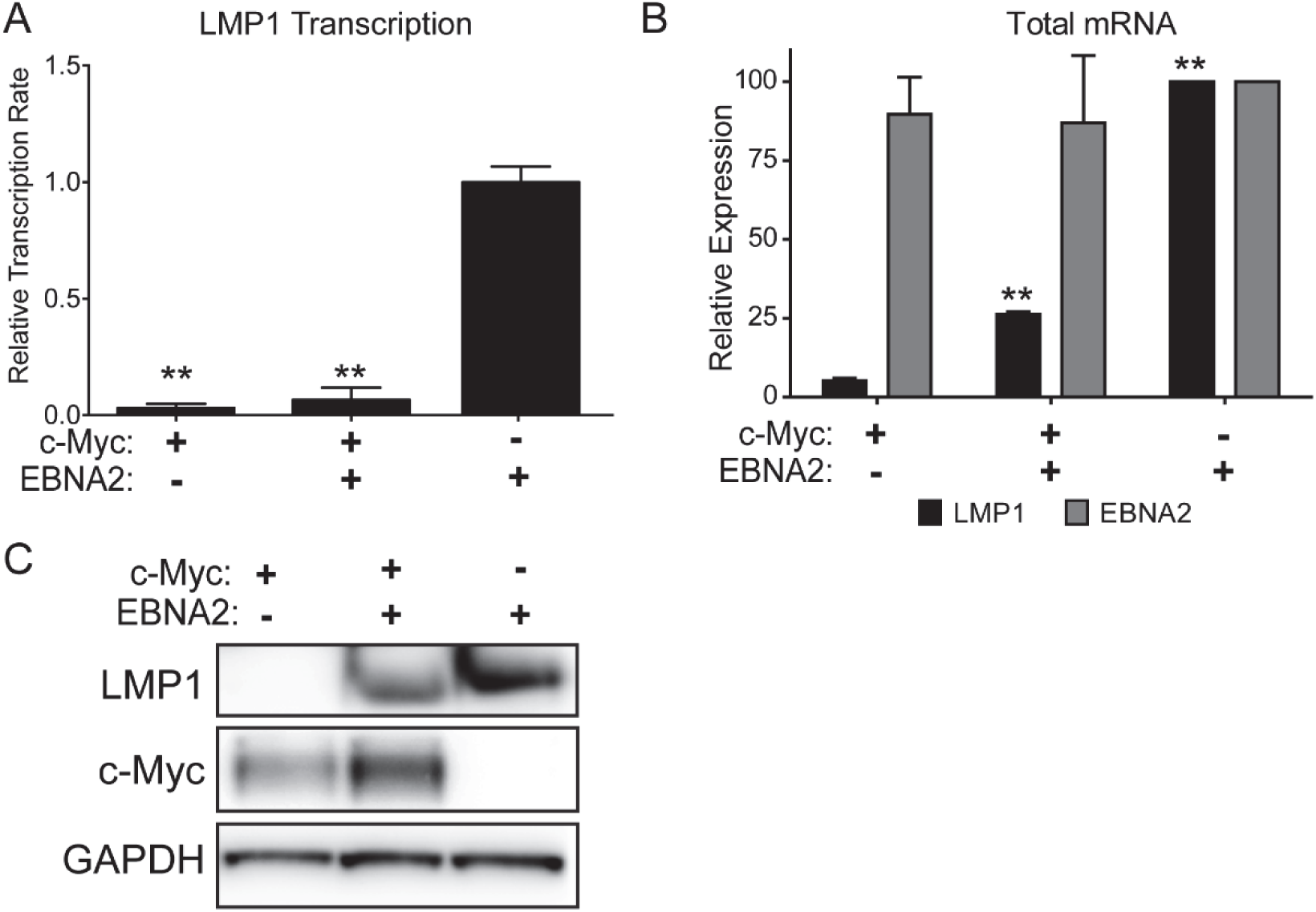
P493-6 system reveals c-Myc inhibits LMP1 transcription and protein expression. (A) RT-qPCR analysis of LMP1 mRNA transcription rate from 4sU metabolic labeling experiments in P493-6 cells in the indicated transcriptional states. All values are normalized to the “LCL” state.(B) RT-qPCR analysis for total LMP1 and EBNA2 mRNA from 4sU metabolic labeling experiments in P493-6 cells in the indicated transcriptional states. All values are normalized to the “LCL” state.(C) Western Blot analysis for LMP1 and c-Myc protein of the P493-6 cells grown in the indicated transcriptional states. For all panels, each bar is representative of at least 3 independent experiments. All error bars denote SEM. ^∗∗^ denotes p<0.01 by one tailed student’s pairwise t-test.

### c-Myc suppresses LMP1 expression early after primary B-cell infection

To test the hypothesis that c-Myc is responsible for LMP1 suppression early after infection, we chose to target c-Myc pharmacologically given the intractability of reverse genetic experiments in primary human B cells. The expression of c-Myc is exquisitely sensitivity to inhibition of bromodomain and extra-terminal motif (BET) domain containing transcriptional activators including Brd4 (43, 44). Therefore, we used two independent BET inhibitors, JQ1 and OTX015, to suppress c-Myc and assay LMP1 transcription in EBV-infected cells. First, we treated LCLs with the BET inhibitors and found that c-Myc mRNA levels decreased and LMP1 mRNA levels increased in a dose-dependent manner (**Fig. 5A-B**). We next treated EBV-infected PBMCs 7 days post infection with BET inhibitors. Consistently, we observed a dose-dependent decrease in c-Myc mRNA and increase in LMP1 mRNA following both JQ1 and OTX015 treatment (**Fig. 5C-D**). Therefore, c-Myc activity negatively correlates with LMP1 expression both early and late after infection.

**Figure 5.**
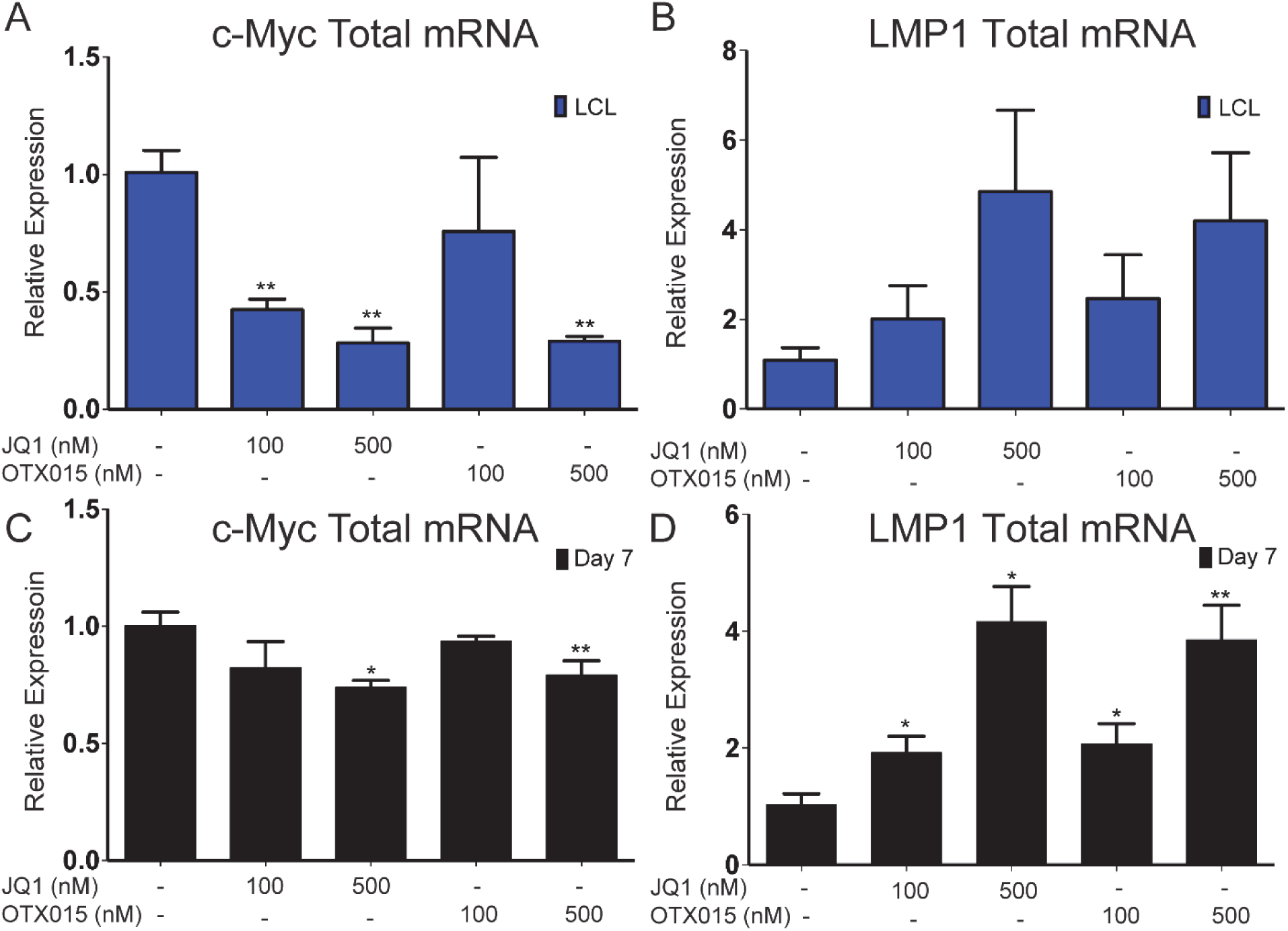
BET Inhibition results in a dose dependent increase in LMP1 and simultaneous decrease in c-Myc mRNA in both primary EBV infected B cells and LCLs. (A) LCLs were treated with the indicated concentrations of either JQ1 or OTX015 or 0.1% DMSO for 24 hours before total mRNA was harvested for RT-qPCR analysis for c-Myc. (B) Similar experiments as in (A) but RT-qPCR was performed for LMP1. (C) Peripheral blood mononuclear cells (PBMCs) were isolated from whole blood and infected with EBV. On day 7, they were treated with the indicated concentrations of JQ1 or OTX015 or 0.1% DMSO for 24 hours before total mRNA was collected for RT-qPCR analysis for c-Myc. (D) Similar experiments as in (C) but RT-qPCR was performed for LMP1. For all panels, each bar represents the average of 3 independent donors. All error bars denote SEM. ^∗^ denotes p<0.05 and ^∗∗^ denotes p<0.01 by one tailed student’s pairwise t-test.

## DISCUSSION

In this study, we determined that transcription of the EBV major latent oncoprotein, LMP1, is suppressed by c-Myc early after primary B cell infection. Transcription of LMP1 increased twenty-five-fold from day 7 to LCL, while the mRNA half-life increased two-fold, leading to an overall fifty-fold increase in LMP1 expression. This change correlated with an approximate two-fold increase in the activating histone mark, H3K9Ac, at the LMP1 promoter, but the occupancy of the major LMP1 trans-activator EBNA2, as well as CTCF-binding to the CTCF-RE, did not change over the same time period. Furthermore, cellular c-Myc was shown to negatively regulate LMP1 transcription and total LMP1 RNA levels in both the setting of highly expressed endogenous c-Myc in a Burkitt lymphoma cell line as well as direct c-Myc overexpression in an LCL. Finally, inactivating c-Myc expression with either of the BRD4-inhibiting compounds, JQ1 or OTX015, led to a dose dependent increase in LMP1 expression both in LCLs and early after primary infection of B cells.

The small but significant change in LMP1 half-life could be explained as a post-transcriptional effect mediated by miRNAs that are known to target the LMP1 3’UTR. Previous work has shown that the miR17~92 family of c-Myc-induced miRNAs target LMP1 and removal of this miRNA family increased LMP1 protein production (27). Furthermore, our group has shown that miR17~92 are expressed at the highest levels early after infection when LMP1 mRNA and mRNA half-life is lowest (45). This low level “fine-tuning” of LMP1 expression might be critical in the immortalized state when high levels of LMP1 can lead to cytostatic effects on cell growth (28, 30).

While the LMP1 promoter has been extensively studied in the immortalized LCL, no one has previously characterized promoter occupancy early after infection. We have shown that early after infection EBNA2 is recruited normally to its response element in the LMP1 promoter as well as several canonical cellular targets including CD23 and c-Myc. This implies that there must be a repressive element present in the LMP1p that supersedes the E2RE’s function at early times after infection. Such an element has been characterized in Burkitt lymphoma cell lines, and it is responsive to members of the Myc family including MAD1 and MAX (46). Alternatively, c-Myc activation could lead to decreased LMP1p occupancy and transcription through less direct mechanisms. Previously c-Myc signaling and NFκB signaling have been shown to activate opposing and mutually exclusive growth programs (34). In this way, direct actions of c-Myc or indirect actions of Myc-induced target genes might lead to less active NFκB transcriptional subunits that can no longer auto-regulate and act in a feed-forward manner on the LMP1p (20).

Many questions still remain regarding the functional relevance for EBV in delaying LMP1 expression. While still important for survival at late stages during infection, it has been shown that EBV can protect B cells from apoptosis in the absence of NFκB activity by activating the anti-apoptotic cellular MCL-1 protein using EBNA3A (36). Additionally, our group has shown that exogenously activating NFκB during the early stage after infection when LMP1 is low increases the transformation rate of the virus (35). However, if the ability of c-Myc to downregulate LMP1 expression was detrimental, one would assume that the virus would evolve to lack such constraints.

Alternatively, we propose c-Myc repression of LMP1 may be beneficial in the life cycle of EBV to maintain low levels of LMP1-induced NFκB early after infection. It has been shown that c-Myc overexpression suppresses recognition of EBV-infected cells by CD8+ T cells (47, 48). In contrast, NFκB activity is known to enhance MHC-mediated antigen presentation, making LCLs excellent targets for CD8+ T cell killing (49). Thus, NFκB activation in EBV-infected B cells in an immune-competent host might ride a fine line between survival and immune recognition. Given our data, we propose that early EBV-infected B cells with high c-Myc and low NFκB activity provide an ideal setting to escape CD8+ T-cell recognition. This is particularly important in seropositive individuals where reactivation and *de novo* naïve B-cell infection is thought to occur in the setting of a robust CD8+ T-cell response. The re-seeding of the latency reservoir from these naïve-infected cells into the memory B-cell compartment would then proceed with attenuated MHC presentation of viral antigens. Therefore, c-Myc-suppressed LMP1/NFκB activity together with the recently described EBV miRNA-mediated attenuation of T-cell recognition and killing (50, 51) are likely the key elements of latent EBV-mediated immune evasion in the immune-competent host.

## MATERIALS AND METHODS

### Cell lines, culture conditions, and viruses

Buffy coats were obtained from normal human donors through the Gulf Coast Regional Blood Center (Houston, TX) and peripheral blood mononuclear cells (PBMCs) were isolated by Ficoll Histopaque-1077 gradient (Sigma, H8889). B95-8 strain of Epstein-Barr virus was produced from the B95-8 Z-HT cell line as previously described (52). Virus infections were performed in bulk by adding 50 μL of filtered B95-8 supernatant to 1×10^6^ PBMCs.

Cell lines were cultured in RPMI 1640 media supplemented with 10-15% heat inactivated fetal bovine serum (Corning), 2 mM L-Glutamine, 100 U/ml penicillin, 100 μg/ml streptomycin (Invitrogen), and 0.5 μg/mL Cyclosporine A (Sigma). P493-6 cells (a kind gift of Dr. Georg Bornkamm, Helmholtz Zentrum München) were cultured with 10% tetracycline-free FBS (Hyclone SH30070), 1 μM β-Estradiol, and 1 μg/mL Tetracycline. All cells were cultured at 37°C in a humidified incubator at 5% CO_2_.

### Flow cytometry and sorting

To track proliferation, cells were stained with CellTrace Violet (Invitrogen, C34557), a fluorescent proliferation-tracking dye. Cells were first washed in FACS buffer (5% FBS in PBS), stained with the appropriate antibody for 30min-1hr at 4°C in the dark, and then washed again before being analyzed on a BD FACS Canto II.

Proliferating infected B cells were sorted to a pure population of CD19^+^/CellTraceViolet^lo^ on a MoFlo Astrios Cell Sorter at the Duke Cancer Institute Flow Cytometry Shared Resource. Mouse anti-human CD19 antibody (clone 33-6-6; gift from Tom Tedder, Duke University Medical School) conjugated with either APC or PE was used as a surface B cell marker in flow cytometry.

### Nascent RNA extraction and profiling

To assess both mRNA transcription and half-life cells were treated with 4-thiouridine (4sU, Sigma) (200 μM) for exactly one hour. Upon harvesting, total RNA was extracted via TRIzol following the manufacturer’s protocol (Life Technologies). 4sU-labelled nascent RNA was then biotinylated using a highly efficient crosslinking reaction using EZ-Link Biotin-HPDP (Pierce) dissolved in DMSO at a concentration of 1 mg/mL or MTSEA Biotin-XX (Biotium Cat# 90066) at 20 ng/uL dissolved in DMSO, and labeled RNA was separated from the total population using streptavidin MyOne C1 Dynabeads (Invitrogen) as previously described (39). Subsequently, three populations of RNA were reverse transcribed into cDNA using the High Capacity cDNA kit (Applied Biosystems): total RNA (T), unlabeled RNA (U), and nascent RNA (N). Quantitative real-time PCR was then performed on these three populations to discover total RNA abundance, relative transcription rates, and mRNA half-life as previously described (53). In brief, the abundance of mRNA was represented by the measurement of the total RNA sample (*T*). Relative transcription was represented by the labeled fraction of mRNA (*N*). The decay rate (*DR*) was calculated from measurements of nascent (*N*) and unlabeled (*U*) mRNA, as a function of *N/U*-ln(1 – *N/U*). An apparent RNA half-life was calculated using the decay rate, –*t* × [ln(2)/*DR*], where *t* is the time of 4sU incorporation (1 h for the purposes of these experiments). Two assumptions of this method are that transcription and stability are constant over the period of measurement.

### Chromatin Immunoprecipitation

Chromatin Immunoprecipitation was performed using ChIP-IT High Sensitivity kit using the manufactures directions (Active Motif). DNA was sonicated for 45 min with 30 second on/off cycles on a Bioruptor (Diagenode). ChIP antibodies include EBNA2 (PE2; gift from Elliot Kieff), H3K9ac (Active Motif cat # 51252 Clone 1B10) and CTCF (Active Motif cat #61311). Quantitative real-time PCR primer sets include the C promoter (F: CCTAGGCCAGCCAGAGATAAT, R: AGATAGCACTCGACGCACTG), LMP1 promoter (F: GGCCAAGTGCAACAGGAA, R: GCAGATTACACTGCCGCTTC), c-Myc Enhancer 525 kb upstream (F: CTAGTAGCAGGTGATGGGTTATG, R: CCTTTGGACCAGAAGAGGATG), c-Myc Enhancer 432 kb upstream (F: ACAGCCAGAGGTATTGGAAC, R: GGAAGGAACGAAACCCTAGAA), CD23 promoter (F: GATCGGCCATAGTGGTATGATT, R: CTCAGGTAAGAGAATTGGGTGAG), CTCF-RE (F: CCACTAGGAACCCAAGATCAA, R: GCCCGCTTCTTCGTATATGT), and EBER/OriP (F: GGGAAATGAGGGTTAGCATAGG, R: CAAGTCTACATCTCCTCAAGACAG. As a negative control for the EBV genome EBNA3C (F: CAAGGTGCATTTACCCCACTG and R: GGGCAGGTCCGTGAGAACT) was used. As negative controls form the human genome two regions Neg1 (F: CCAATAACAGAAGCATTAAAATTCA, R: TTCAAGCACAGGCATACAGG) and Neg2 (F: TCTCTGGGGAGATGGATTACA, R: CGTGAATCCTTTATTCTTGGAA) were used.

### Gene Expression Analysis

Total RNA was isolated from cells by using a Qiagen RNeasy kit and then reverse transcribed to generate cDNA with the High Capacity cDNA kit (Applied Biosystems). Quantitative PCR was performed by using SYBR green (Quanta Biosciences) in an Applied Biosystems Step One Plus instrument. Primer sets include c-Myc (F: CTCCATGAGGAGACACCGC, R: GAGCCTGCCTCTTTTCCACA), LMP1 (F: AATTTGCACGGACAGGCATT, R: AAGGCCAAAAGCTGCCAGAT), EBNA2 (F: GCTTAGCCAGTAACCCAGCACT, R: TGCTTAGAAGGTT GTTGGCATG), GAPDH (F: TGCACCACCAACT GCTTAGC, R: GGCATGGACTGTGGTCATGAG) and SETDB1 (F: TCCATGGCATGGTGGAGCGG, R: GAGAGGGTTCTTGCCCCGGT).

### Western blot

Cells were pelleted and washed in PBS, and then lysed in 0.1% Triton-containing buffer with Complete protease inhibitors. All protein lysates were run on NuPage 4–12% gradient gels (LifeTechnology) and transferred to PVDF membrane (GE Healthcare). Membranes were blocked in 5% milk in TBST and stained with primary antibody overnight at +4°C, followed by a wash and staining with secondary HRP-conjugated antibody for 1 hour at room temperature. Antibodies include LMP1 (S12; gift from Elliot Kieff, Harvard Medical School), c-Myc (Santa Cruz Biotechnology, SC-764), and GAPDH (BioChain Institute, #Y3322).

### BRD4 Inhibition

(+)-JQ1 and OTX015 were purchased from Selleckchem (cat. #S7110 and #S7360, respectively).

#### For LCLs

LCLs were seeded at 3^∗^10^^^5/mL in the presence of either 0.1% DMSO, 100 or 500 nM JQ1 or 100 or 500 nM OTX015 for 24 hours. After 24 hours total mRNA was harvested using the Qiagen RNeasy mini kit (ref #74104) according to manufacturer’s instructions. 1 microgram of harvest mRNA was reverse transcribed into cDNA using Applied Biosystems High-Capacity cDNA reverse transcription kit (cat #4368814) according to manufacturer’s instructions. RT-qPCR analysis was performed using SYBR Green detection based system (Quanta Bio cat #95072-05K) with 5 ng of cDNA used per reaction using an Applied Biosystems StepOne Plus Real Time PCR System. All values were normalized to the endogenous loading control SETDB1. Relative expression values were calculated using the ΔΔCT method.

#### For PBMCs

Peripheral Blood Mononuclear Cells (PBMCs) were isolated from whole blood, infected with EBV B95-8 and seeded at 1^∗^10^^^6/mL. Seven days post infection the cells were treated with either 0.1% DMSO, 100 or 500 nM JQ1 or 100 or 500 nM OTX015. After 24 hours, total mRNA was harvested as previously described. 1 μg of total RNA was reverse transcribed into cDNA as previously described. RT-qPCR analysis was conducted as previously described.

## ACKNOWLEDGMENTS

We thank Lynn Martinek, Nancy Martin, and Mike Cook for extensive help in flow-based cytometry experiments. Special thanks are due to Jack Keene and Jeff Blackinton for assistance with 4sU labeling experiments. This work was supported by National Institutes of Health (NIH) Grants R01-CA140337 and R01-DE025994 (to M.A.L.), F31-CA180451 (to A.M.P), and T32-CA009111 (to A.M.P. and J.E.M). Additional funding came from the Duke CFAR, an NIH funded program, 5P30-AI064518, and an American Cancer Society grant RSG-13-228-01-MPC (Both to M.A.L.).

